# Loss of Syndecan-3 Drives a Pro-inflammatory, EndMT-prone Phenotype in Aortic Valve Endothelial Cells

**DOI:** 10.64898/2026.09.03.749289

**Authors:** George I. Ezeokeke, Guy C. Bedford, Joseph J.P. Stroschein, Anael Roig-Gicquel, Isaac B. Hilton, K. Jane Grande-Allen

**Affiliations:** Department of Bioengineering, Rice University, Houston, TX 77030, USA; Department of Biosciences, Rice University, Houston, TX 77030, USA

**Author notes:** **Address for correspondence:** K. Jane Grande-Allen, PhD, Department of Bioengineering, Rice University 6100 Main Street, MS-142, Houston, TX 77030, USA Email: [ ].

**Keywords:** calcific aortic valve disease, syndecan-3, endothelial glycocalyx, NF-κB, endothelial-to-mesenchymal transition, valve endothelial cells

## Abstract

The earliest events in calcific aortic valve disease (CAVD) occur at the valve endothelium, where the cells lining the leaflet undergo NF-κB (Nuclear Factor kappa B)–driven inflammatory activation and endothelial-to-mesenchymal transition (EndMT). The endothelial glycocalyx (EG) sets the threshold at which endothelial cells (ECs) respond to such cues, yet its composition and function in the aortic valve remain almost entirely uncharacterized. Here we define the cell-surface proteoglycan (CPG) landscape of human aortic valve endothelial cells (HAVECs) under inflammatory stimulation and identify syndecan-3 (SDC3) as a regulator of valve endothelial activation. Among the proteoglycans examined, SDC3 was highly expressed and inflammation-responsive. Silencing SDC3 in unstimulated HAVECs was sufficient to upregulate NF-κB pathway components and the leukocyte adhesion molecules vascular cell adhesion molecule-1 (VCAM1), intercellular adhesion molecule-1 (ICAM1), and E-selectin, and it amplified their induction upon TNF-α challenge. Pharmacological inhibition of IKKβ abolished this difference, placing SDC3 upstream of NF-κB, while SDC3 overexpression attenuated tumor necrosis factor-alpha (TNF-α)–induced VCAM1, the inverse of the knockdown effect. Loss of SDC3 further primed HAVECs toward a mesenchymal, EndMT-prone phenotype under transforming growth factor-beta 2 (TGF-β2) stimulation, marked by increased smooth muscle protein 22-alpha (SM22α) and reduced platelet endothelial cell adhesion molecule (PECAM1/CD31). Together, these findings identify SDC3 as an endogenous restraint on the inflammatory and mesenchymal programs that characterize the endothelial phase of CAVD and establish the valve endothelial glycocalyx as a previously unappreciated yet potent regulator of aortic valve endothelial dysfunction. Preserving or augmenting SDC3 may warrant investigation as a means of holding valve endothelial cells in a quiescent state.

## INTRODUCTION

Calcific aortic valve disease (CAVD) is a slow and progressive pathology of the aortic valve that ranges from mild valve thickening, often termed aortic sclerosis, to severe calcification with impaired leaflet motion, termed aortic stenosis [1]. Upon the emergence of symptoms, CAVD is associated with high morbidity and mortality, contributing to over 100,000 deaths globally each year, with a worldwide prevalence that continues to rise and currently affects an estimated 9.6 to 12.6 million people, rendering it an increasingly urgent public health burden [2]. The only definitive treatments remain surgical or transcatheter aortic valve replacement [3–4], which restore leaflet function and improve survival but do not address the biological processes that drive the disease.

Because the underlying pathology is left untreated, these interventions manage the mechanical endpoint rather than halt or reverse the disease. Implanted bioprosthetic valves are themselves subject to structural degeneration that limits their lifespan, so that younger patients in particular face the prospect of repeated valve reintervention over their lifetime [3]. The absence of disease-modifying treatment reflects a longstanding and incomplete understanding of the cellular and molecular mechanisms that initiate and drive valve calcification [3].

The aortic valve leaflet is a thin, layered connective tissue structure populated by two principal cell types. Aortic valve interstitial cells (VICs) reside within the leaflet interior [1] and maintain the extracellular matrix that gives the leaflet its mechanical integrity, while a single layer of aortic valve endothelial cells (ECs) covers the leaflet surface [1], forming the interface between the tissue and the flowing blood. These two populations occupy distinct microenvironments and are exposed to different mechanical and biochemical cues, and both have been implicated in the disease process. Most mechanistic studies have focused on the VICs [5], the predominant cell population within the leaflet, which undergo osteogenic differentiation and directly deposit the calcific nodules that define end-stage disease [6–7]. However, calcification does not arise in isolation; the earliest pathological events in CAVD occur at the valve surface, where the aortic valve ECs that form the leaflet’s protective lining become dysfunctional in response to disturbed hemodynamics and inflammatory stimuli [8]. Yet relative to the VICs, the biology of the valve endothelium remains comparatively underexplored [5]. This is a critical oversight, as valvular endothelial dysfunction influences a multi-pronged pathological program that actively drives disease progression. Dysfunctional aortic valve ECs orchestrate an array of destructive processes. They initiate immune cell infiltration by upregulating vascular cell adhesion molecule-1 (VCAM1), intercellular adhesion molecule-1 (ICAM1), and E-selectin to recruit circulating leukocytes and fuel chronic inflammation [9], and they undergo endothelial-to-mesenchymal transition (EndMT), shedding their endothelial identity to acquire a mesenchymal, VIC-like phenotype that contributes to CAVD progression [9,10]. Ultimately, this cellular reprogramming positions the endothelium as an important contributor to both CAVD initiation and downstream progression.

Projecting from the luminal membrane of every EC is the endothelial glycocalyx (EG), a dense, carbohydrate-rich layer of proteoglycans, glycosaminoglycans, and associated glycoproteins that constitutes the cell’s primary interface with the circulating environment [11]. Far from being a passive structural layer, the glycocalyx transduces hemodynamic shear into intracellular signaling, regulates vascular permeability, and sequesters cytokines and growth factors at the cell surface, thereby setting the threshold at which an endothelial cell responds to inflammatory and mechanical cues [12]. EG degradation is an early and consequential event in vascular disease, preceding and precipitating overt endothelial dysfunction [13–14]. Among its key constituents are cell surface heparan sulfate proteoglycans, comprising the transmembrane syndecans (SDC) and the glypicans (GPC), which are tethered to the cell membrane by a glycosylphosphatidylinositol anchor. Both families serve as signaling co-receptors linking extracellular ligand engagement to intracellular pathways [11]. Yet despite the glycocalyx’s recognized importance in vascular biology, characterization of its composition and function in the aortic valve endothelium has not advanced beyond the earliest ultrastructural descriptions of the valve surface coat decades ago [15–16], leaving how this protective layer governs valve endothelial behavior largely unknown.

Within the SDC family, SDC3 is the least studied member [17–18], and its biology has historically been mischaracterized. Early studies, identifying its abundant expression in the developing and adult nervous system, largely restricted SDC3 to a neuronal role [19–20], and only more recently has it been recognized as a broader participant in inflammatory signaling and angiogenesis across non-neuronal tissues [21]. Whether SDC3 contributes to the inflammatory regulation of the valve endothelium, as it does in other non-neuronal tissues, is not yet understood.

Here, we provide the first systematic characterization of the cell-surface proteoglycan (CPG) landscape of aortic valve endothelial cells under baseline conditions as well as inflammatory insult, and within it identify SDC3 as a previously unrecognized regulator of aortic valve endothelial activation. We show that SDC3 influences NF-κB signaling in aortic valve ECs, such that its loss alone is sufficient to raise the expression of the canonical activation markers VCAM-1, ICAM-1, and E-selectin, and to prime the cells for amplified responses to inflammatory challenge. The loss of SDC3 further predisposes aortic valve ECs to EndMT, linking a single glycocalyx proteoglycan to both programs that define the endothelial dysfunction phase of CAVD. In identifying this functional constituent of the valve EG, these findings establish this layer as a previously unappreciated regulator of aortic valve endothelial dysfunction and identify SDC3 as a point of entry for interrogating, and potentially modifying, disease at its endothelial origin.

## METHODS

### Cell Culture & Growth Factor Stimulation

Human Aortic Valve Endothelial Cells (HAVECs) and Human Coronary Artery Endothelial Cells (HCAECs) were obtained from Lonza and maintained in complete endothelial growth medium (EGM-2) (Lonza, cat. 3162) supplemented with 5% fetal bovine serum (FBS) (Fisher Scientific, cat.

MT35010CV) and penicillin–streptomycin (Millipore Sigma, cat. P078) in a humidified incubator with 5% CO₂ at 37°C. Both cell types were used within passages 4–6 for all experiments. Prior to treatment with recombinant human tumor necrosis factor alpha (TNF-α) (R&D Systems, cat. 210-TA), lipopolysaccharide (LPS; 100 ng/mL) (MilliporeSigma, cat. L4391), or recombinant human transforming growth factor-beta 2 (TGF-β2; 10 ng/mL) (R&D Systems, cat. 302-B2), cells were serum-starved overnight in endothelial basal medium (EBM-2) (Lonza, cat. 3156) supplemented with 2% FBS. TNF-α was used at the concentrations (100 ng/mL or 10 ng/mL) indicated in the corresponding figure legends. All treatments were performed in the same 2% FBS EBM-2 medium.

### Western Blot

HAVECs and HCAECs cultured in 6 cm dishes or 6-well plates were used for Western blot analysis. Cells were washed with ice-cold PBS and lysed on ice in RIPA buffer (ThermoFisher, cat. 89901) supplemented with protease inhibitor (ThermoFisher, cat. 87785). Lysates were cleared by centrifugation, and total protein concentration was determined by BCA assay (ThermoFisher, cat. PI23227). Equal amounts of protein were denatured in Laemmli sample buffer and separated by SDS-PAGE using polyacrylamide gels. Proteins were transferred to PVDF membranes, blocked with EveryBlot Blocking Buffer (Bio-Rad, cat. 12010020), and incubated with indicated primary antibodies overnight at 4 °C. Membranes were then washed in TBST, incubated with HRP-conjugated secondary antibodies, washed again, and developed using chemiluminescent substrate.

Immunoreactive bands were quantified using Image Lab software (Bio-Rad). The following primary antibodies were used: anti-SDC3 (1:500) (Abcam, cat. 155952), anti-SDC1 (1:1000) (Proteintech, cat. 10593-1-AP), anti-VCAM1 (1:1000) (ThermoFisher, cat. MA5-31965), anti-E-selectin/SELE (1:1000) (Cell Signaling Technology, cat. 61280), anti-FLAG (1:1000) (Millipore Sigma, cat. F1804), anti-SM22α/transgelin (1:1000) (Cell Signaling Technology, cat. 36090), anti-CD31 (1:300) (ThermoFisher, cat. BS-0468R), anti-GAPDH (1:5000) (Proteintech, cat. HRP-60024), and anti-β-tubulin (1:1000) (ThermoFisher, cat. MA5-16308-HRP) as loading controls. HRP-conjugated secondary antibodies against rabbit IgG (ThermoFisher, cat. 31460) or mouse IgG (ThermoFisher, cat. 31430) were used at 1:5000 as appropriate for the host species of each primary antibody.

### SDC3 Silencing in HAVECs

HAVECs were seeded onto 24-well plates or 6-well plates and transfected at approximately 80% confluency using Opti-MEM (ThermoFisher, cat. 31985070) and Lipofectamine RNAiMAX (ThermoFisher, cat. 13778030). Transfections were performed in a total volume of 1 mL Opti-MEM containing 0.6 μL Lipofectamine RNAiMAX per well in 24-well plates, or 2 mL Opti-MEM containing 2.5 μL Lipofectamine RNAiMAX per well in 6-well plates. For SDC3 knockdown, a custom 27mer SDC3 siRNA duplex (Integrated DNA Technologies) was used at 40 nM (sense: 5′-rGrCrCrCrUrUrGrArArArUrCrArUrCrUrArGrArCrArCrUGC-3′; antisense: 5′-rGrCrArGrUrGrUrCrUrArGrArUrGrArUrUrUrCrArArGrGrGrCrUrG-3′). A Trilencer-27 Universal scrambled negative control siRNA duplex (Origene, cat. SR30004) was used as the non-targeting control (NTC) at the same concentration. Twenty-four hours after transfection, the transfection medium was replaced with full EGM-2 supplemented with 5% FBS. Cells were serum-starved in EBM-2 supplemented with 2% FBS for 6 h before growth factor stimulation.

### mRNA Synthesis & Transfection for SDC3 Overexpression

3xFLAG-tagged syndecan-3 (3xFLAG-SDC3) mRNA was produced by in vitro transcription (IVT). The transcription template was a T7-promoter plasmid built with the Takara Cloning Kit for mRNA Template (cat. 6143), in which a 3xFLAG tag was inserted immediately downstream of the start codon of the full-length human SDC3 coding sequence by NEBuilder HiFi DNA assembly (New England Biolabs). The assembly was transformed into Stable3 competent cells, selected on kanamycin (50 µg/ml), and confirmed by whole-plasmid sequencing (Quintara Biosciences) before use. The template carries a plasmid-encoded poly(A) tract, so no enzymatic polyadenylation was performed. The sequence-verified plasmid was linearized with HindIII-HF (New England Biolabs, cat. R3104S), which cuts immediately 3’ of the poly(A) tract, and column-purified before use. IVT was performed with the TriLink CleanCap AG (3’ OMe) CleanScript IVT Kit (cat. K-7413) according to the manufacturer’s specifications, with N1-methylpseudouridine substituted for UTP, typically in 20 µl reactions, which yielded sufficient mRNA for experiments. mRNA was purified by lithium chloride precipitation (2.5 M final concentration), washed in 70% ethanol, resuspended in nuclease-free water, and stored at −80 °C. Product integrity (approximately 1,670 nt) was verified on a 1% agarose gel, after denaturing at 70 °C for 10 min, with NEB RNA Loading Dye (2X, cat. B0363S) and an ssRNA ladder (New England Biolabs, cat. N0362S). HAVECs were transfected at approximately 80% confluency with SDC3 mRNA using Opti-MEM (ThermoFisher, cat. 31985070) and Lipofectamine MessengerMAX (ThermoFisher, cat. LMRNA). SDC3 mRNA was used at 1 μg/mL, unless otherwise indicated, and at 0.005, 0.025, 0.1, 0.5, and 1 μg/mL for dose-response experiments. Control cells were transfected with Lipofectamine MessengerMAX alone.

### RNA isolation, cDNA synthesis, and quantitative reverse transcription PCR

Cells were lysed in TRIzol (ThermoFisher, cat. 15596026), and total RNA was extracted using the Direct-zol™ RNA Microprep kit (Zymo Research, cat. R2062) according to the manufacturer’s instructions. Briefly, lysates were mixed with an equal volume of 95% ethanol and transferred to a Zymo-Spin™ IC Column in a collection tube. Samples were treated with DNase I for 15 min at room temperature, washed with RNA wash buffer, and eluted in 15 μL DNase/RNase-free water. RNA purity (A260/280 > 1.8) was assessed using a NanoDrop 2000 spectrophotometer, and RNA was stored at −80 °C until further processing. Complementary DNA (cDNA) was synthesized from purified RNA using the High-Capacity cDNA Reverse Transcription Kit (Applied Biosystems, cat. 4368813) according to the manufacturer’s instructions and stored at −20 °C until gene expression analysis. RT-qPCR was performed using iTaq™ Universal Probes Supermix (Bio-Rad, cat. 1725134) according to the manufacturer’s instructions, with the following TaqMan Gene Expression Assays (Applied Biosystems): *SDC1* (Hs04966523_m1), *SDC2* (Hs01081432_m1), *SDC3* (Hs01568665_m1), *SDC4* (Hs01120908_m1), *GPC1* (Hs00892476_m1), *VCAM1* (Hs01003372_m1), *ICAM1* (Hs00164932_m1), *SELE* (Hs00174057_m1), *RELA* (Hs01042014_m1), *NFKB1* (Hs00765730_m1), *NFKBIA* (Hs00355671_g1), *TNFAIP3* (Hs00234713_m1), *ACTA2* (Hs00426835_g1), *TAGLN* (Hs01038777_g1), *CLDN5* (Hs00533949_s1), *KDR* (Hs00911700_m1), *SMAD7* (Hs00998193_m1), *PMEPA1* (Hs00375306_m1), and *GAPDH* (Hs02786624_g1) as the endogenous control. Relative gene expression was calculated using the 2^−ΔΔCt method, normalized to GAPDH and expressed relative to the indicated control condition.

### Inhibition of NF-κB Signaling with BMS-345541

HAVECs were transfected with non-targeting control (NTC) or SDC3-targeting siRNA at 40 nM in 24-well plates as described above. At 72 hours post-transfection, cells were pretreated with the selective allosteric IKKβ/α inhibitor BMS-345541 hydrochloride (Sigma-Aldrich, cat. B9935) or a matched dimethyl sulfoxide (DMSO) vehicle control. BMS-345541 was reconstituted in sterile DMSO to a stock concentration of 11.43 mM and used at a final working concentration of 11.43 μM (0.1% DMSO final) in EBM-2 supplemented with 2% FBS. Following a 1-hour pretreatment at 37 °C, the pretreatment medium was aspirated and replaced with fresh EBM-2 supplemented with 2% FBS, containing 11.43 μM BMS-345541 (or matched DMSO vehicle) and 100 ng/mL TNF-α to maintain sustained allosteric inhibition throughout the treatment period. Cells were incubated for an additional 4 hours, then harvested for RNA.

### Statistical Analysis

Statistical analysis was performed using GraphPad Prism software version 10 (GraphPad Software Inc.). Data are presented as mean ± standard deviation (SD), with n indicating the number of independent biological replicates as specified in each figure legend. Comparisons between two conditions were analyzed using an unpaired two-tailed Student’s t-test. Comparisons across multiple groups were analyzed using one-way or two-way analysis of variance (ANOVA), as appropriate to the experimental design, followed by Tukey’s post hoc test for multiple comparisons. Statistical significance was set at p < 0.05, and significance levels are denoted in the figures as *p < 0.05, **p < 0.01, ***p < 0.001, and ****p < 0.0001.

## RESULTS

### A. SDC3 is highly expressed in aortic valve endothelial cells and is upregulated by inflammatory stimulation

Primary HAVECs were cultured *in vitro*, and the expression levels (relative to GAPDH) of the four SDCs (*SDC1*, *SDC2*, *SDC3*, and *SDC4*) and *GPC1* were determined by RT-qPCR. GPC1 was expressed at the highest level, followed by *SDC3* and *SDC4*, while *SDC1* and *SDC2* were expressed at low levels (Figure 1A). This pattern closely paralleled that reported in human umbilical vein endothelial cells (HUVECs), in which *SDC3* and *SDC4* are the predominant syndecans and *SDC1* and *SDC2* are expressed at low levels [22]. To address the possible importance of CPGs in the inflammatory responses of HAVECs, we cultured HAVECs in the presence of two major inflammatory mediators, LPS and TNF-α, and determined mRNA expression relative to untreated controls. As shown in Figure 1B, individual CPGs responded differently to inflammatory stimulation: *SDC2* and *GPC1* were progressively downregulated by TNF-α, each reaching approximately 0.3–0.4-fold by 48 h, whereas *SDC3* and *SDC4* were upregulated. *SDC4* showed the greatest fold induction, reaching approximately 20-fold under TNF-α and 12-fold under LPS at 4 h, whereas *SDC3* rose more modestly, reaching approximately 1.4-fold under both LPS and TNF-α by 48 h. To determine whether the CPG responses were specific to the HAVECs, we performed the same experiments in human coronary artery endothelial cells (HCAECs) as a vascular endothelial control, quantifying mRNA expression across a time course of 4, 24, and 48 h. Of the CPGs examined, SDC1 was notable for its cell type-dependent behavior. In HAVECs, *SDC1* mRNA was unchanged at 4 h but was downregulated by TNF-α to approximately 0.5-fold at 24 h and 0.6-fold at 48 h, while LPS modestly increased SDC1 to approximately 1.4-fold by 48 h (Figure 1C). In HCAECs, *SDC1* mRNA showed a biphasic response to TNF-α, decreasing at 4 h, recovering by 24 h, and decreasing again at 48 h. The clearest distinction between HAVECs and HCAECs emerged at the protein level. Following 48 h of stimulation, SDC1 protein was markedly decreased by TNF-α in HAVECs but increased by both LPS and TNF-α in HCAECs (Figure 1D).

**Figure 1.**
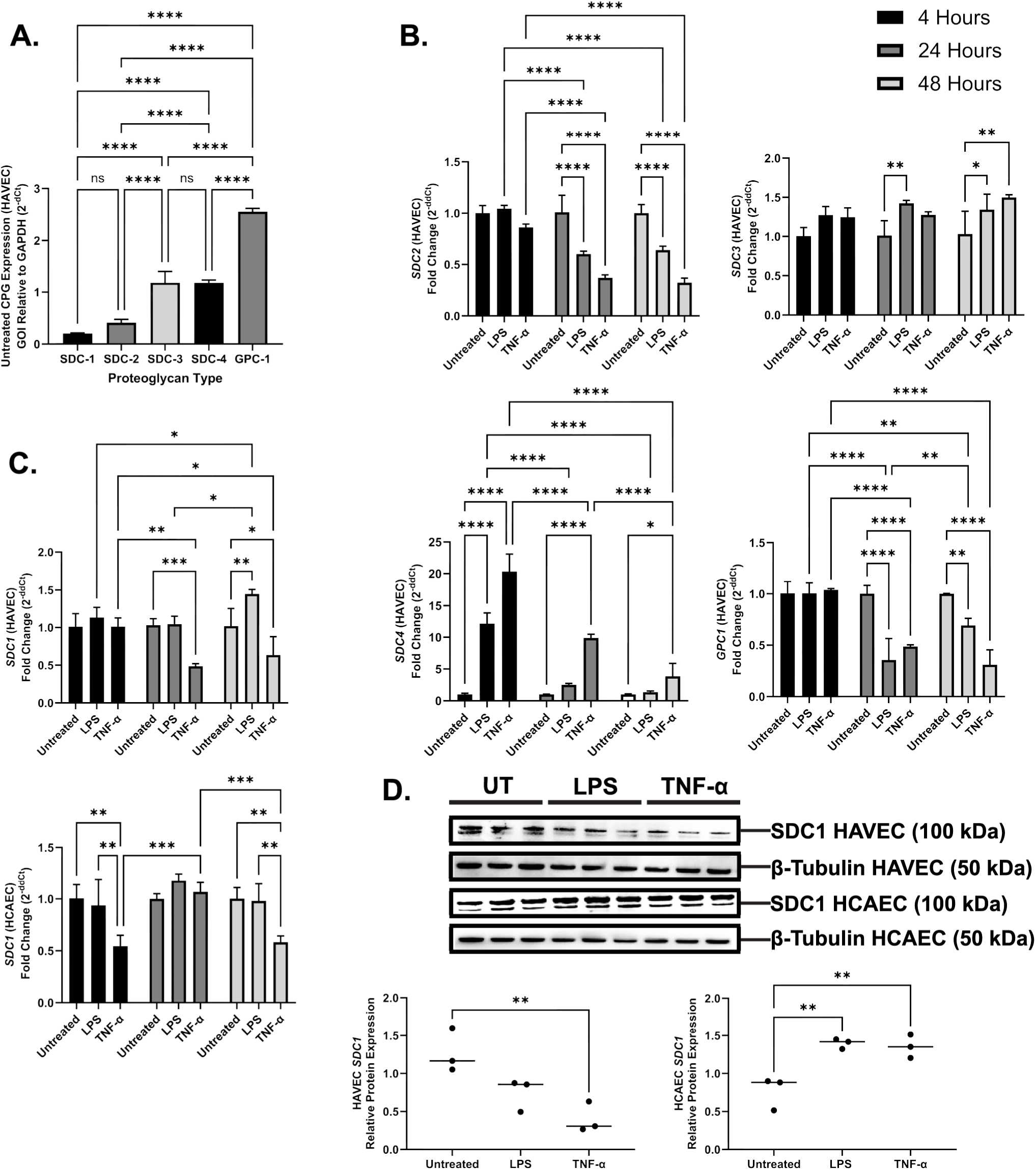
Characterization of cell-surface proteoglycan expression and its inflammatory regulation in human aortic valve endothelial cells. (**A**) Baseline expression of cell-surface proteoglycans (CPGs) in unstimulated human aortic valve endothelial cells (HAVECs), quantified by RT-qPCR relative to GAPDH (n=3). (**B**) RT-qPCR quantification of SDC-2, SDC-3, SDC-4, and GPC-1 transcripts in HAVECs following stimulation with LPS (100 ng/mL) or TNF-α (100 ng/mL) for 4, 24, or 48 hours, expressed as fold change relative to untreated control (n=3). (**C**) RT-qPCR quantification of SDC-1 transcript expression in HAVECs and human coronary artery endothelial cells (HCAECs) under identical stimulation conditions (n=3). (**D**) Representative Western blots and corresponding densitometric quantification of SDC-1 protein in HAVECs and HCAECs following 48 h of LPS (100 ng/mL) or TNF-α (100 ng/mL) treatment, normalized to β-tubulin. Data are presented as mean ± SD (n = 3 per condition); *p < 0.05, **p < 0.01, ***p < 0.001, ****p < 0.0001; ns, not significant.

Although SDC4 demonstrated the most robust response, further characterization offers limited novelty given its well-established role in endothelial inflammatory signaling [22–24]. Prior work has demonstrated that SDC3 exerts tissue-selective, vascular bed dependent functions, acting as a pro-inflammatory or anti-inflammatory mediator depending on the tissue [25]. Whether SDC3 serves a proinflammatory or anti-inflammatory function within the specialized microenvironment of the aortic valve endothelium remains entirely unknown. Given its high baseline expression and inflammatory-mediated inducibility in HAVECs, we prioritized SDC3 in subsequent studies to define its role in aortic valvular endothelial inflammation.

### B. Loss of SDC3 alone, without inflammatory insult, is sufficient to upregulate NF-κB pathway components and leukocyte adhesion molecules

Having identified SDC3 as an inflammation-responsive CPG in HAVECs, we next asked whether SDC3 itself regulates the inflammatory state of these cells. We silenced SDC3 in HAVECs using siRNA and confirmed efficient knockdown at both the transcript and protein levels relative to a non-targeting control (NTC), with *SDC3* transcript reduced by approximately 90% (to approximately 0.1-fold of NTC) (Figure 2A). To determine the consequences of SDC3 loss, we first examined the leukocyte adhesion molecules that mediate endothelial recruitment of circulating immune cells. In the absence of any inflammatory stimulus, SDC3-silenced HAVECs significantly increased the expression of *VCAM1*, *ICAM1*, and E-selectin (*SELE*) relative to NTC, reaching approximately 2.9-, 1.8-, and 3.3-fold, respectively (Figure 2B). Because these adhesion molecules are transcriptional targets of NF-κB signaling [26], we next examined the expression of core NF-κB pathway components. SDC3 knockdown significantly upregulated *NFKBIA*, *NFKB1*, and *RELA*, to approximately 1.7-, 1.5-, and 1.3-fold, respectively (Figure 2C), indicating that the loss of SDC3 elevates NF-κB pathway expression even in the resting, unstimulated state.

**Figure 2.**
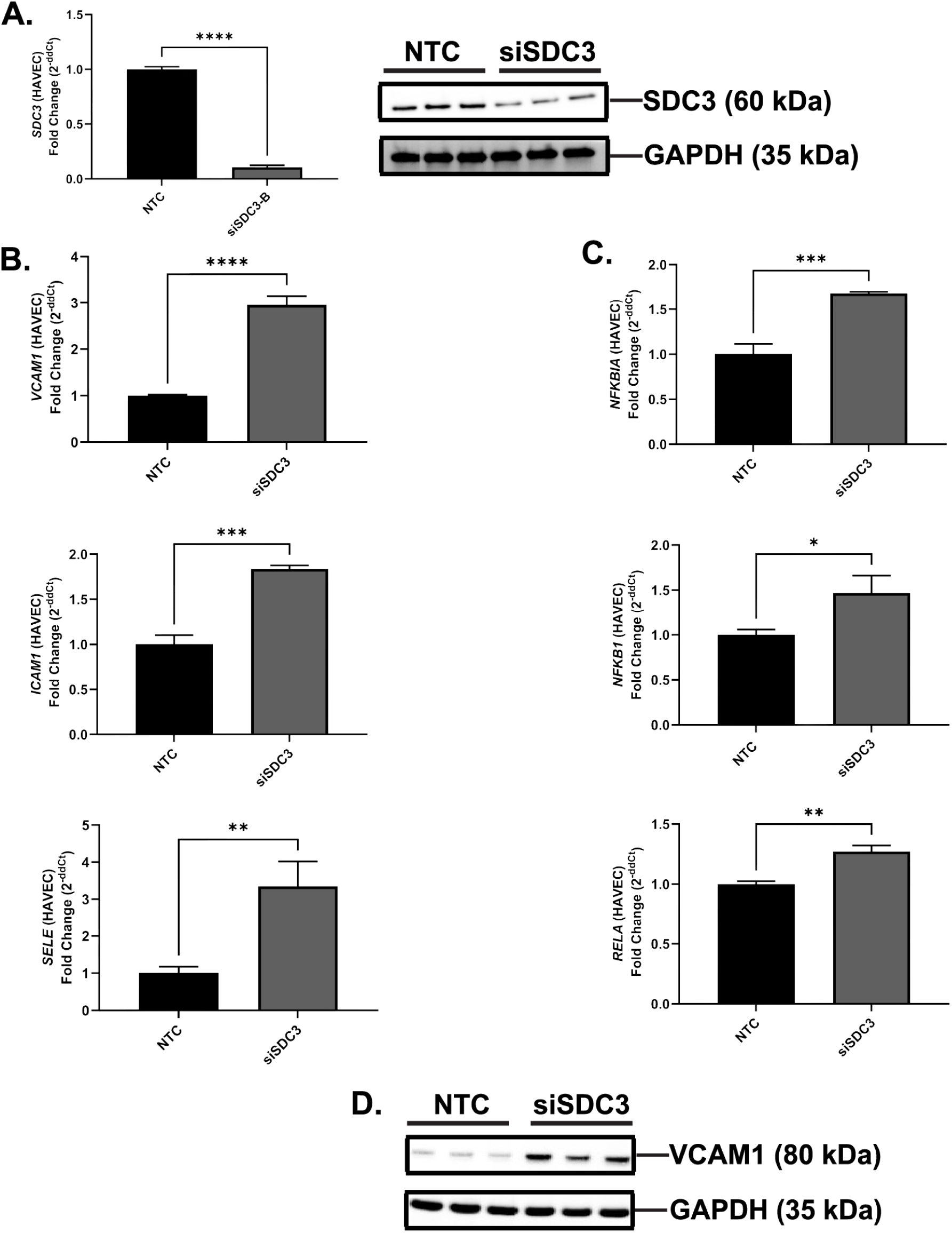
Loss of *SDC3* alone, without inflammatory insult, is sufficient to upregulate NF-κB pathway components and leukocyte adhesion molecules in human aortic valve endothelial cells. **(A)** *SDC3* transcript expression by RT-qPCR (left) and representative Western blot of SDC3 protein with GAPDH loading control (right) in NTC and siSDC3 HAVECs. **(B)** RT-qPCR quantification of *VCAM1*, *ICAM1*, and E-selectin (*SELE*) in NTC and siSDC3 HAVECs. **(C)** RT-qPCR quantification of *NFKBIA*, *NFKB1*, and *RELA* in NTC and siSDC3 HAVECs. **(D)** Representative Western blot of VCAM1 protein with GAPDH loading control in NTC and siSDC3 HAVECs. Data are presented as mean ± SD (n = 3 per condition); *p < 0.05, **p < 0.01, ***p < 0.001, ****p < 0.0001.

To confirm that this transcriptional upregulation was reflected at the protein level, we assessed VCAM1 protein in unstimulated NTC and siSDC3 HAVECs. Consistent with the mRNA data, VCAM1 protein was elevated following SDC3 knockdown (Figure 2D). Together, these results demonstrate that SDC3 loss alone, in the absence of any inflammatory insult, is sufficient to upregulate NF-κB pathway components and their downstream leukocyte adhesion molecule targets, identifying SDC3 as an endogenous restraint on the inflammatory tone of the valve endothelium.

### C. Loss of SDC3 primes aortic valve endothelial cells for an amplified inflammatory response to TNF-α

Having established that SDC3 loss elevates inflammatory tone in resting HAVECs, we next asked whether SDC3 also shapes the magnitude of the response to an inflammatory challenge. NTC and siSDC3 HAVECs were stimulated with TNF-α for 4 h, and adhesion molecule and NF-κB pathway expression were compared to untreated controls.

TNF-α strongly induced the leukocyte adhesion molecules VCAM1, ICAM1, and E-selectin (SELE) in NTC cells. This induction was significantly greater in SDC3-silenced cells: siSDC3 HAVECs stimulated with TNF-α expressed higher levels of all three adhesion molecules than TNF-α– stimulated NTC cells, reaching approximately 149-versus 89-fold for *VCAM1*, 101-versus 65-fold for *ICAM1*, and 58,000-versus 11,700-fold for *SELE* (Figure 3A). A comparable pattern was observed for the NF-κB pathway components *NFKBIA*, *NFKB1*, and *RELA*, each of which was more strongly induced by TNF-α in siSDC3 cells than in NTC controls, reaching approximately 36-versus 15-fold, 20-versus 10.5-fold, and 1.8-versus 1.6-fold, respectively (Figure 3B). To determine whether this amplified response extended to the protein level, we examined VCAM1 and E-selectin protein across a time course of TNF-α stimulation (4, 8, and 16 h), chosen to capture the distinct induction kinetics of the two adhesion molecules, with E-selectin induced early and VCAM1 more sustained [27]. Both proteins were more strongly induced in siSDC3 HAVECs than in NTC cells, with E-Selectin peaking at 8 h and VCAM1 at 16 h (Figure 3C), confirming that the potentiated response is reflected in protein abundance.

**Figure 3.**
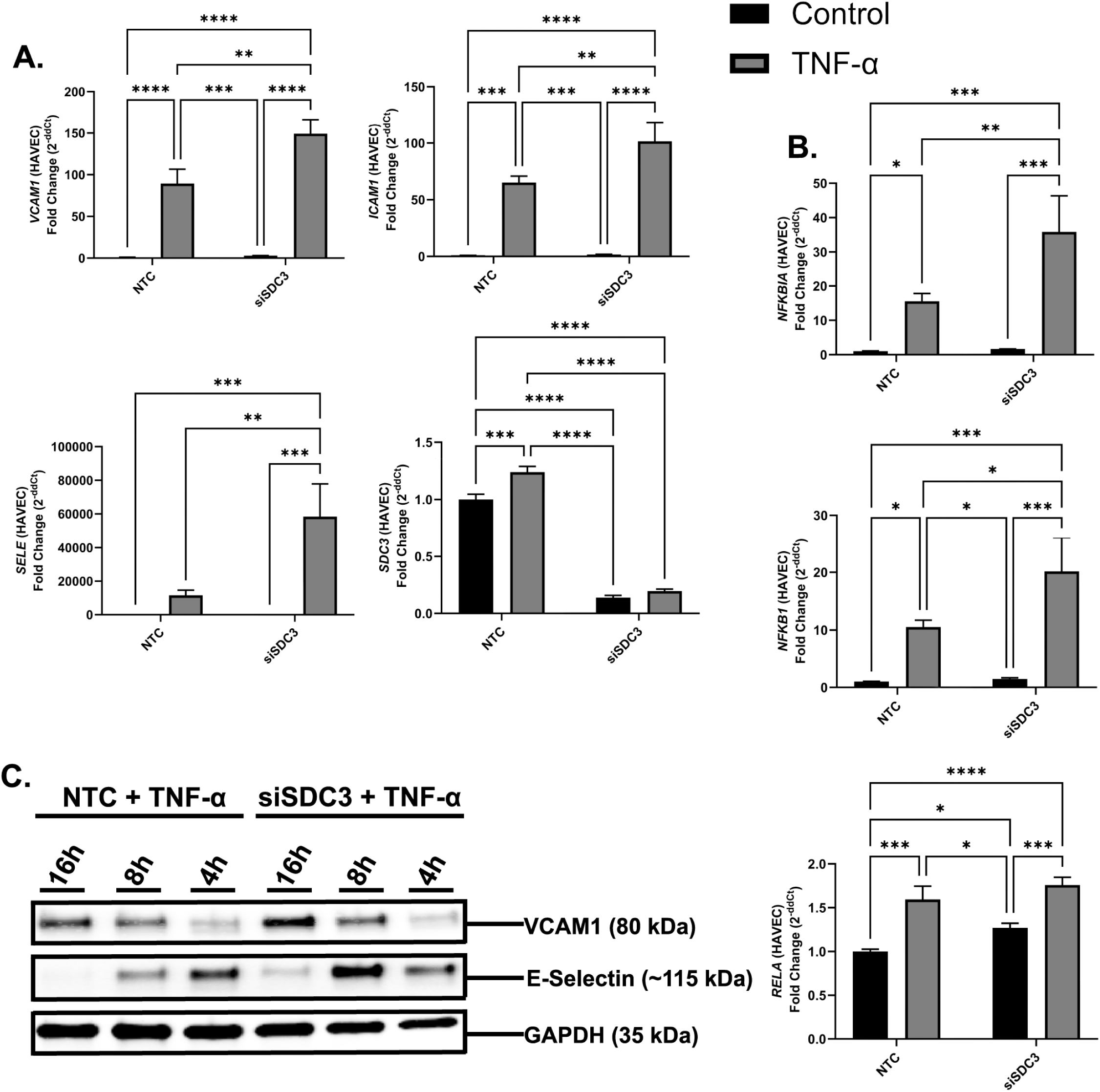
Loss of *SDC3* primes aortic valve endothelial cells for an amplified inflammatory response to TNF-α. **(A)** RT-qPCR quantification of *VCAM1*, *ICAM1*, and E-selectin (*SELE*), and of *SDC3*, in NTC and siSDC-3 HAVECs, with and without TNF-α (100 ng/mL) stimulation for 4 h. **(B)** RT-qPCR quantification of *NFKBIA*, *NFKB1*, and *RELA* in NTC and siSDC-3 HAVECs, with and without TNF-α (100 ng/mL) stimulation for 4 h. **(C)** Representative Western blot of VCAM1 and E-selectin protein across a TNF-α (10 ng/mL) stimulation time course (4, 8, 16 h) in NTC and siSDC-3 HAVECs, with GAPDH as loading control. All qPCR data are expressed as fold change relative to NTC (2^−ddCt); *p < 0.05, **p < 0.01, ***p < 0.001, ****p < 0.0001.

### D. Inhibitor of nuclear factor kappa-B kinase subunit β (IKKβ) inhibition and SDC3 overexpression suggest that SDC3 restrains TNF-α–induced inflammatory activation of HAVECs upstream of NF-κB

The preceding results indicated that SDC3 loss elevates gene expression of key NF-κB pathway components and potentiates the TNF-α response but did not establish whether these effects depend on NF-κB signaling. To test this, we stimulated NTC and siSDC3 HAVECs with TNF-α in the presence of either DMSO vehicle or IKKβ inhibitor BMS-345541 (BMS).

For these inhibitor experiments, fold changes were normalized to the NTC vehicle + TNF-α condition, set to 1.0, rather than to unstimulated cells as in the preceding figures. In vehicle-treated cells, SDC3 knockdown reproduced the priming phenotype: siSDC3 HAVECs expressed significantly higher levels of *VCAM1*, *ICAM1*, and E-selectin (*SELE*) than NTC cells following TNF-α stimulation, reaching approximately 2.6-versus 1.0-fold, 1.8-versus 1.0-fold, and 6.0-versus 1.3-fold, respectively (Figure 4A–C). IKKβ inhibition abolished this difference. In BMS-treated cells, adhesion molecule expression was strongly suppressed to near-baseline in both genotypes, and the elevated expression seen in siSDC3 cells was no longer present, such that siSDC3 and NTC cells were statistically indistinguishable under BMS (Figure 4A–C). These results indicate that the priming produced by SDC3 loss requires IKKβ signaling. In addition, BMS treatment reduced *SDC3* expression in NTC cells by approximately half relative to vehicle (from approximately 1.0- to 0.5-fold; Figure 4D), indicating that SDC3 expression itself may be dependent on IKKβ activity.

**Figure 4.**
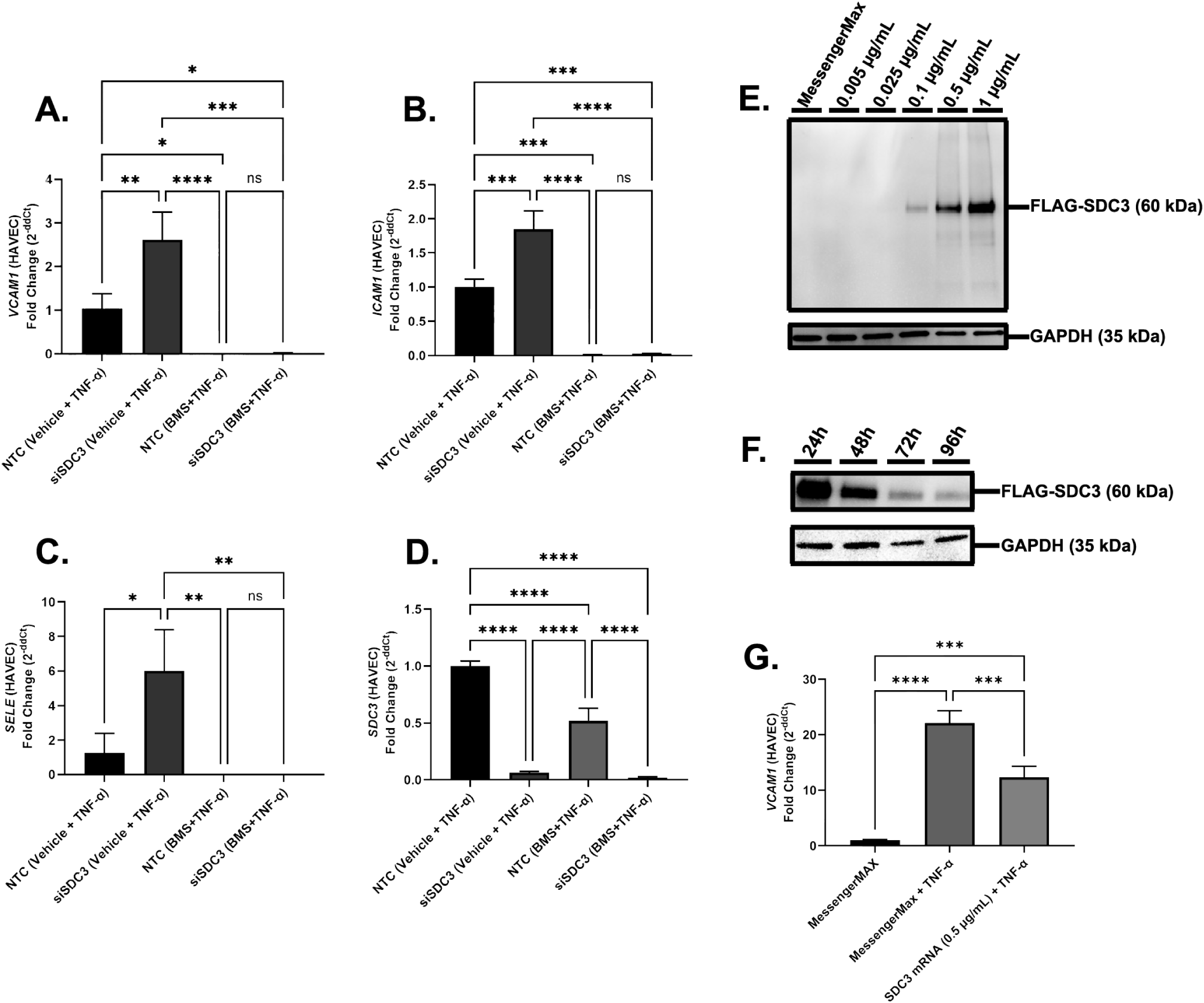
IKKβ inhibition and syndecan-3 overexpression confirm that SDC3 restrains the TNF-α response upstream of NF-κB. (A–C) RT-qPCR quantification of *VCAM1* (A), *ICAM1* **(B)**, and E-selectin (*SELE*) **(C)** in NTC and siSDC3 HAVECs stimulated with TNF-α (100 ng/mL) for 4 h in the presence of DMSO vehicle or the IKKβ inhibitor BMS-345541 (BMS). **(D)** RT-qPCR quantification of *SDC3* across the same conditions. **(E)** Representative Western blot of FLAG-tagged SDC3 protein following transfection of HAVECs with increasing concentrations of SDC3 in vitro–transcribed (IVT) mRNA (0.005–1 µg/mL) or MessengerMAX reagent alone, with GAPDH as loading control. **(F)** Representative Western blot of FLAG-SDC3 protein at 24, 48, 72, and 96 h following transfection of SDC3 IVT mRNA, with GAPDH as loading control. **(G)** RT-qPCR quantification of *VCAM1* in HAVECs transfected with MessengerMAX alone, MessengerMAX with TNF-α (100 ng/mL) for 4 h, or SDC3 IVT mRNA (0.5 µg/mL) with TNF-α (100 ng/mL) for 4 h; *p < 0.05, **p < 0.01, ***p < 0.001, ****p < 0.0001; ns, not significant.

To complement these loss-of-function experiments, we asked whether increasing SDC3 expression would produce the opposite effect. We generated an in vitro–transcribed (IVT) mRNA encoding FLAG-tagged SDC3 and introduced it into HAVECs. Transfection produced a dose-dependent increase in SDC3 protein across a range of mRNA concentrations (Figure 4E), and a time-course analysis showed that SDC3 protein peaked at 24 h and declined progressively through 96 h, defining the window of overexpression (Figure 4F). Using these parameters, we tested the effect of SDC3 overexpression on TNF-α–induced *VCAM1*. TNF-α strongly induced *VCAM1* in control (MessengerMAX-transfected) cells, reaching approximately 22-fold, and this induction was significantly attenuated to approximately 12-fold in cells overexpressing SDC3 (Figure 4G). Thus, increasing SDC3 expression suppressed the TNF-α–driven induction of *VCAM1*, the inverse of the effect produced by SDC3 knockdown.

### E. Loss of SDC3 primes aortic valve endothelial cells for a mesenchymal, EndMT-prone phenotype

NF-κB and TGF-β signaling have been shown to act synergistically to drive EndMT, such that inflammatory activators of the NF-κB pathway markedly potentiate TGF-β–induced transition; this cooperativity has been demonstrated for potent NF-κB activators including TNF-α and IL-1β [28–29]. For the TGF-β arm of this study, TGF-β2 was selected, as it is the most potent inducer of EndMT among the TGF-β isoforms in endothelial cells, with TGF-β1 and TGF-β3 relying in part on TGF-β2 to trigger the transition [30]. We therefore reasoned that, because SDC3 loss elevated NF-κB–driven adhesion molecule expression in HAVECs even at baseline (Figure 2), the same loss might also predispose these cells to EndMT, either by initiating the transition or by priming them to undergo it more readily upon TGF-β2 stimulation.

To test this, we transfected HAVECs with NTC or SDC3-targeting siRNA for 48 h and stimulated them with TNF-α, TGF-β2, or both for 48 h, and quantified EndMT-associated transcripts and proteins. Interestingly, *SDC3* expression was significantly induced in NTC HAVECs under combined TNF-α + TGF-β2 stimulation, rising approximately 1.9-fold over control (Figure 5A). This paralleled the TNF-α–driven induction of *SDC3* we observed in Figures 1 and 3, where inflammatory stimulation of HAVECs upregulated *SDC3* and where its loss primed HAVECs for an amplified response to TNF-α.

**Figure 5.**
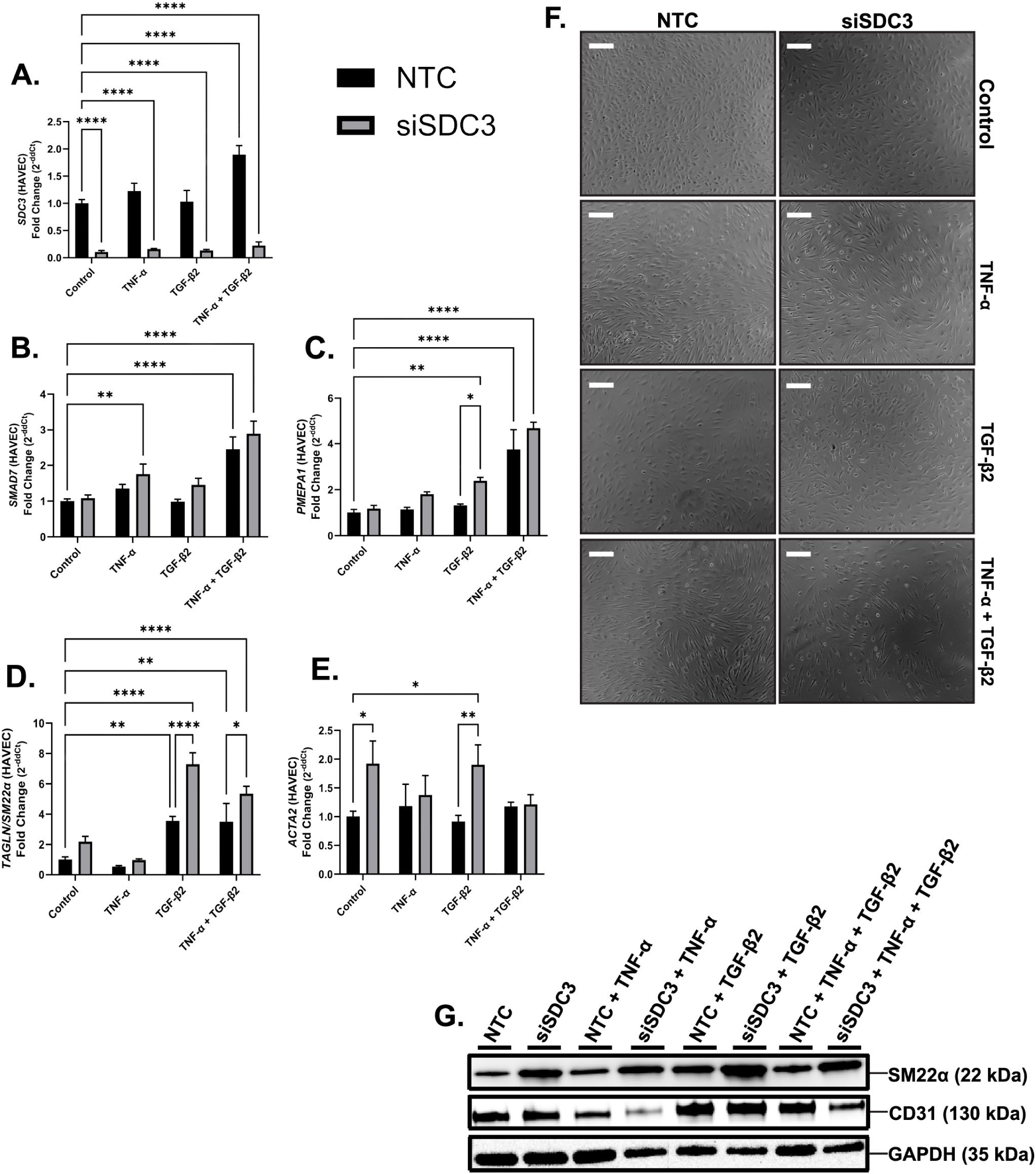
Loss of SDC3 primes aortic valve endothelial cells for a mesenchymal, EndMT-prone phenotype under TGF-β2 stimulation. (A–E) RT-qPCR quantification of *SDC3* (A), *SMAD7* (B), *PMEPA1* (C), *TAGLN*/SM22α (D), and *ACTA2* (E) in NTC and siSDC3 HAVECs under control conditions or following stimulation with TNF-α (10 ng/mL), TGF-β2 (10 ng/mL), or both for 48 h. Data are expressed as fold change relative to NTC control (2^−ddCt) (n=3). **(F)** Representative phase-contrast images of NTC and siSDC3 HAVECs under the same conditions. Scale bars, [100] µm. **(G)** Representative Western blot of SM22α and CD31 protein in NTC and siSDC3 HAVECs across the four conditions, with GAPDH as loading control; *p < 0.05, **p < 0.01, ***p < 0.001, ****p < 0.0001.

SMAD7 is an inhibitory SMAD that acts as a negative feedback regulator of TGF-β signaling and whose transcription is induced by TGF-β [31]. Beyond this canonical role, SMAD7 also participates in crosstalk with the NF-κB pathway, restraining NF-κB activation by inhibiting IκB degradation [32], which places it at the intersection of the two pathways central to this study. *SMAD7* expression was elevated in siSDC3 cells relative to NTC in every condition examined (Figure 5B), indicating a consistent, genotype-associated increase in the feedback response, although this difference did not reach significance in all groups. Notably, TNF-α significantly induced *SMAD7* in siSDC3 cells (approximately 1.8-fold relative to untreated control), whereas the corresponding induction in NTC cells (approximately 1.4-fold) did not reach significance. Because this condition contains no TGF-β2, the *SMAD7* induction in SDC3-deficient cells is most consistent with their heightened NF-κB tone driving its expression, in line with the reciprocal SMAD7–NF-κB crosstalk noted above. *SMAD7* was also elevated under combined stimulation, reaching approximately 2.9-fold in siSDC3 cells versus 2.5-fold in NTC. Because *SMAD7* mRNA induction tends to peak early [33], we examined an earlier time point and found that at 72 h post-transfection, siSDC3 cells expressed significantly higher *SMAD7* transcripts than NTC cells without stimulation with TNF-α or TGF-β2 (Supplementary Figure 3A).

Prostate transmembrane androgen inducible protein-1 (PMEPA1) is a direct SMAD2/3 transcriptional target that serves as a faithful reporter of canonical TGF-β pathway activity [28]. Under TGF-β2, *PMEPA1* was elevated in siSDC3 cells to approximately 2.4-fold, compared with approximately 1.3-fold in NTC (Figure 5C). Because PMEPA1 reflects active canonical signaling, its elevation suggests that TGF-β/SMAD output was greater in SDC3-deficient cells despite the concurrent rise in SMAD7 expression. This heightened pathway activity was reflected in the mesenchymal effector *TAGLN* (SM22α), which showed the clearest genotype-dependent effect of the panel: under TGF-β2, *TAGLN* reached approximately 7.3-fold in siSDC3 cells versus 3.5-fold in NTC and remained elevated under combined stimulation (approximately 5.4-fold versus 3.5-fold; Figure 5D). The mesenchymal marker *ACTA2* was likewise elevated in siSDC3 cells, most notably under control and TGF-β2 conditions, where siSDC3 cells reached approximately 1.9-fold over the corresponding NTC groups (Figure 5E). Consistent with the loss of endothelial identity that accompanies mesenchymal transition, the endothelial markers *CLDN5* and *KDR* were progressively downregulated under EndMT-inducing stimulation, with both reaching approximately 0.5-fold under combined TNF-α + TGF-β2 relative to unstimulated control (Supplementary Figure 3B, C).

These transcript-level changes were mirrored at the protein level (Figure 5G). SM22α protein was increased in siSDC3 cells relative to NTC across conditions, most prominently under TGF-β2, consistent with the *TAGLN* transcript data, while the endothelial marker CD31 was reduced in siSDC3 cells, most clearly following TNF-α stimulation. The concurrent gain of SM22α and loss of CD31 within the same samples indicates that these opposing changes reflect a genuine phenotypic shift toward a mesenchymal state.

## DISCUSSION

The earliest events in CAVD occur at the endothelial surface of the leaflet, where the lining cells become activated and begin to lose their endothelial identity while gaining a more mesenchymal, VIC-like phenotype. Because these initiating events are endothelial, the factors governing valve endothelial behavior are of particular interest as potential targets for early intervention. In vascular ECs, one such governor is the EG, the carbohydrate-rich surface layer that sets the threshold at which endothelial cells respond to inflammatory and mechanical cues; however, its composition has remained almost entirely uncharacterized in the aortic valve. We reasoned that glycocalyx proteoglycans might regulate the early endothelial activation implicated in valve disease, and we therefore characterized this landscape and defined the function of SDC3 within it. The main findings were that loss of SDC3, in the absence of any inflammatory stimulus, was sufficient to raise NF-κB pathway components and their downstream leukocyte adhesion molecules and amplified the response of these aortic valve ECs to TNF-α. Correspondingly, SDC3 overexpression produced the opposite effect and attenuated TNF-α-induced VCAM1. In addition, loss of SDC3 primed these cells for a mesenchymal, EndMT-prone phenotype upon TGF-β2 stimulation. Together, these results identify SDC3 as an anti-inflammatory cell-surface proteoglycan in aortic valve ECs.

This work also provides, to our knowledge, the first systematic characterization of the EG landscape of aortic valve ECs and its regulation under inflammatory stress, a contribution independent of the identification of SDC3 as an anti-inflammatory proteoglycan within it. This characterization establishes that the valve endothelial glycocalyx is a dynamically and selectively regulated structure, with individual proteoglycans differing in both their baseline abundance and their responses to inflammatory stimulation. Each of the regulated proteoglycans identified here represents a candidate modifier of aortic valve endothelial behavior, and defining their individual functions, or the consequences of coordinated remodeling of the landscape, is an avenue this characterization opens.

Our finding that SDC3 restrains inflammation in the valve endothelium relates to a growing recognition of its role in non-neuronal tissues. SDC3 is the least studied of the syndecans and was long regarded as a predominantly neuronal proteoglycan but has since been shown to influence NF-κB signaling in non-neuronal cells, including a recent demonstration that its overexpression suppresses LPS-induced NF-κB activation in bovine mammary epithelial cells [34]. Its reported effects across tissues nonetheless remain context- and tissue-dependent, with loss of SDC3 proving to be pro-inflammatory in synovial endothelium but, interestingly, anti-inflammatory in dermal and cremaster tissue [25]. Our data place SDC3 in aortic valve ECs in the anti-inflammatory category, consistent with this emerging non-neuronal role, while extending it to the aortic valve endothelium and linking it to EndMT, which prior studies do not address. These vascular-bed-dependent behaviors align with the broader observation that glycocalyx regulation differs across endothelial and tissue types [35], including reports that inflammatory stimulation drives differential glycocalyx shedding between arterial and venous endothelium [36], and with our own observation that SDC1 responded differently to inflammatory stimulation, at both the mRNA and protein levels, in valvular versus coronary (vascular) ECs. A comparable relationship with inflammatory and EndMT-related programs has also been reported for other glycocalyx core proteins. For example, GPC-1 has been shown to protect vascular endothelial cells from stiffness-mediated inflammatory gene expression and EndMT, such that its loss sensitizes them to these programs [37].

An alternative explanation for the amplified inflammatory phenotype of SDC3-silenced HAVECs warrants consideration. Rather than SDC3 acting directly on the NF-κB pathway, its loss could raise inflammatory tone indirectly, for example by altering the expression of other cell-surface proteoglycans, several of which we found to be inflammation-responsive in HAVECs. The IKKβ inhibition experiment constrains this possibility: because blockade of IKKβ eliminated the difference between SDC3-silenced and control HAVECs, abolishing the amplified TNF-α response produced by SDC3 loss, the effect of SDC3 loss requires IKKβ activity and potentially acts at or upstream of the IKK complex rather than downstream of it. This narrows the possibilities but does not fully resolve them, as a compensatory shift in the wider proteoglycan landscape could still act through the same IKKβ-dependent node. Two observations argue against a purely compensatory mechanism. First, the reciprocal effects of SDC3 silencing and overexpression on the same readout are more consistent with a direct action of SDC3 than with an indirect consequence of proteoglycan rebalancing. Second, the same directional relationship between SDC3 and NF-κB has been reported in an unrelated cell type, where SDC3 overexpression suppressed NF-κB activation in bovine mammary epithelial cells [34]; that this relationship holds across multiple cell types with distinct proteoglycan compositions suggests it is an intrinsic property of SDC3 rather than a peculiarity of the valve EG. The further observation that IKKβ inhibition also lowered SDC3 expression in control HAVECs suggests a feedback arrangement, in which NF-κB pathway activity supports SDC3 expression while SDC3 in turn restrains the pathway; this relationship is inferred from expression data and remains to be tested directly.

Beyond its role in inflammation, loss of SDC3 primed aortic valve ECs for EndMT, extending its influence to the second program that defines the endothelial phase of CAVD. The behavior of SMAD7 offers insight into how this occurs. SMAD7, an inhibitory SMAD induced by both TGF-β and NF-κB signaling, was upregulated by TNF-α specifically in SDC3-silenced cells, in keeping with its induction by NF-κB and its reciprocal crosstalk with the pathway [32]. Yet despite this rise in a negative regulator, canonical TGF-β output remained higher in these same cells. We interpret this to mean that SDC3 loss elevates both the transition-driving pathway and its own inhibitory brake, and that the feedback engaged is insufficient to offset the net drive toward transition. This pattern is notable because it indicates that SDC3 loss does not simply add to TGF-β signaling but shifts the balance of a self-regulating circuit through the cells mounting a compensatory SMAD7 response, but that response fails to contain the heightened pathway activity, leaving the cells poised to transition. The convergence of elevated NF-κB tone and unrestrained TGF-β output in SDC3-deficient HAVECs may therefore explain why these cells are primed for EndMT rather than merely more inflamed.

Although this is an in vitro study, the findings carry potential clinical relevance. A CPG that restrains the initiating inflammatory and mesenchymal programs identifies a candidate point of intervention in the early, endothelial phase of CAVD, before the leaflet fibrosis and calcification that define its later stages. The endothelial events characterized here, NF-κB–driven activation and the shift toward a mesenchymal phenotype, occur upstream of the osteogenic differentiation and matrix mineralization that produce the mechanical valve failure of end-stage disease. Because these early endothelial changes precede and help drive that later structural damage, a factor that holds the endothelium in its quiescent state acts at a stage when the process may still be modifiable, before the accumulation of calcific nodules renders it effectively irreversible. Positioning SDC3 and the glycocalyx more broadly at this early node identifies the valve endothelial surface as a compartment worth interrogating for interventions aimed at the origin of the disease rather than at its mechanical endpoint.

This study has several limitations. It was performed entirely in vitro in cultured HAVECs, so determining whether the relationships defined here operate in the intact valve, or are modified by disease in vivo, will require further investigation. Additionally, several of the baseline transcript-level differences between SDC3-silenced and control cells were modest; the more compelling evidence for a phenotypic shift is the concurrent, opposing movement of an endothelial and a mesenchymal marker within the same samples rather than the magnitude of any single change. This distinction matters, as a statistically detectable difference in a low-variability culture system need not correspond to a biologically meaningful one, and whether effects of this magnitude translate to meaningful modification of disease in vivo remains unknown.

The finding that SDC3 restrains NF-κB signaling upstream of IKKβ, yet does so through a mechanism that remains undefined, suggests several directions for further study. The means by which SDC3 restrains IKKβ activity could involve the heparan sulfate chains, the core protein, sequestration of inflammatory ligands at the cell surface, or co-receptor signaling; the IVT mRNA overexpression system established here, which allows structural variants of SDC3 to be introduced and compared, offers one route to distinguishing among these. Because our data derive entirely from cultured cells, immunostaining of SDC3 in stenotic versus non-diseased human aortic valves would test whether the relationships defined in vitro operate in disease, and stable loss-of-function models would allow the inflammatory and mesenchymal consequences of SDC3 loss to be followed over time. Finally, because the glycocalyx is also a principal transducer of hemodynamic shear, and laminar shear stress classically maintains vascular ECs in a quiescent, atheroprotective state [12], whether SDC3 participates in this shear response, and whether its loss compromises the protective phenotype that laminar flow normally sustains, is a further question raised directly by its role as a restraint on endothelial activation.

In conclusion, we have shown SDC3 to be an endogenous restraint on NF-κB-driven inflammation in HAVECs, and its loss primes these cells for the inflammatory activation and mesenchymal transition that characterize the endothelial phase of CAVD. Because increasing SDC3 reverses the inflammatory effect of its loss, preserving or augmenting this proteoglycan, or the glycocalyx more broadly, may warrant investigation as a strategy to keep valve endothelial cells in a quiescent, healthy state.

## ACKNOWLEDGMENTS

Funding was provided by 25DIVSUP1478725 (to G.I.E.).

