## Supplementary Material for "Loss of Syndecan-3 Drives a Pro-inflammatory, EndMT-prone Phenotype in Aortic Valve Endothelial Cells"

**SUPPLEMENTARY FIGURES**

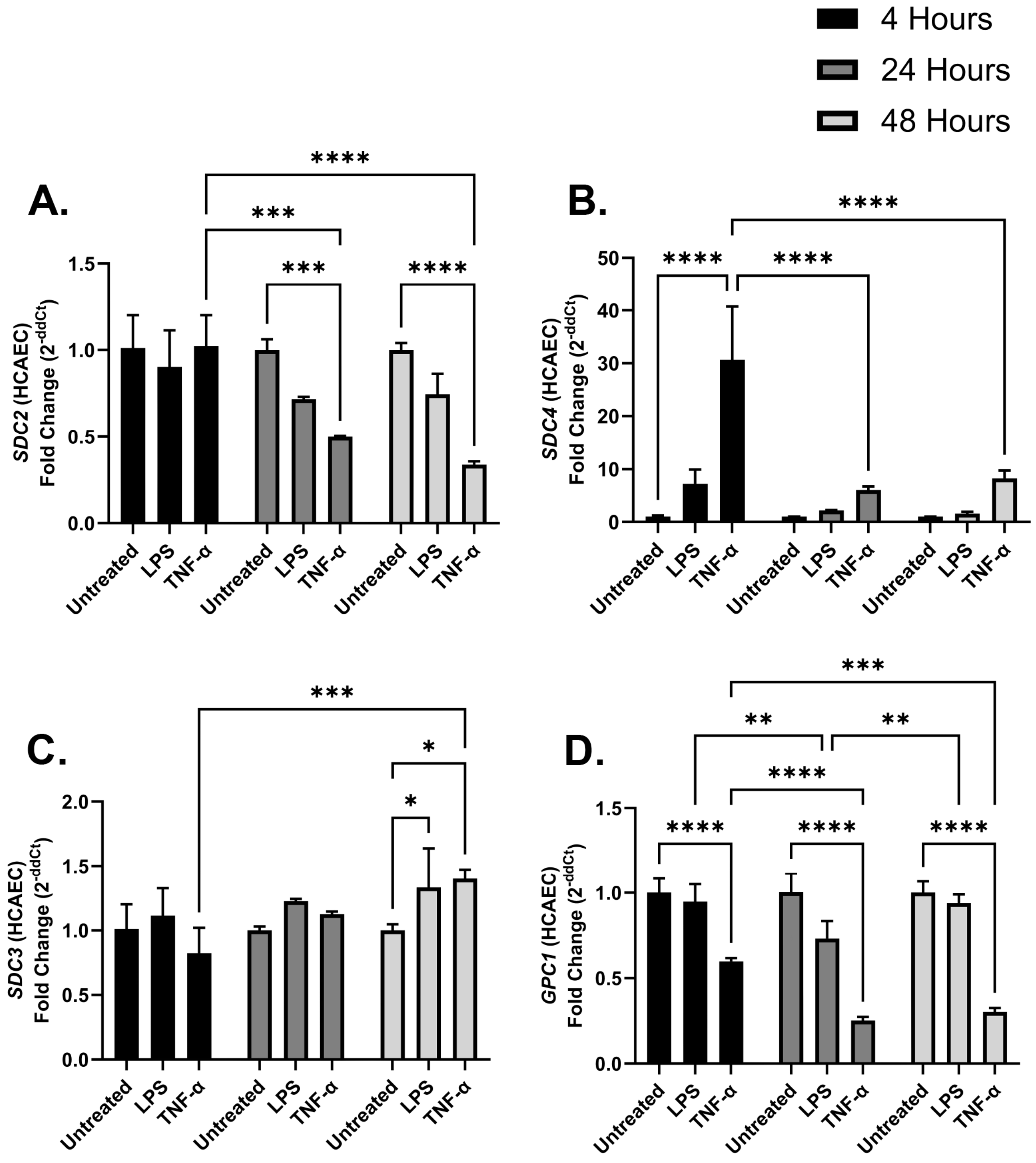


**Supplementary Figure 1. Inflammatory regulation of cell-surface proteoglycans in HCAECs.** RT-qPCR quantification of (A) *SDC2*, (B) *SDC4*, (C) *SDC3*, and (D) *GPC1* transcripts in HCAECs following stimulation with LPS or TNF-α (both at 100 ng/mL) for 4, 24, or 48 h, expressed as fold change relative to untreated control (2^−ΔΔCt). Data are presented as mean ± SD (n = 3 per condition); *p < 0.05, **p < 0.01, ***p < 0.001, ****p < 0.0001.

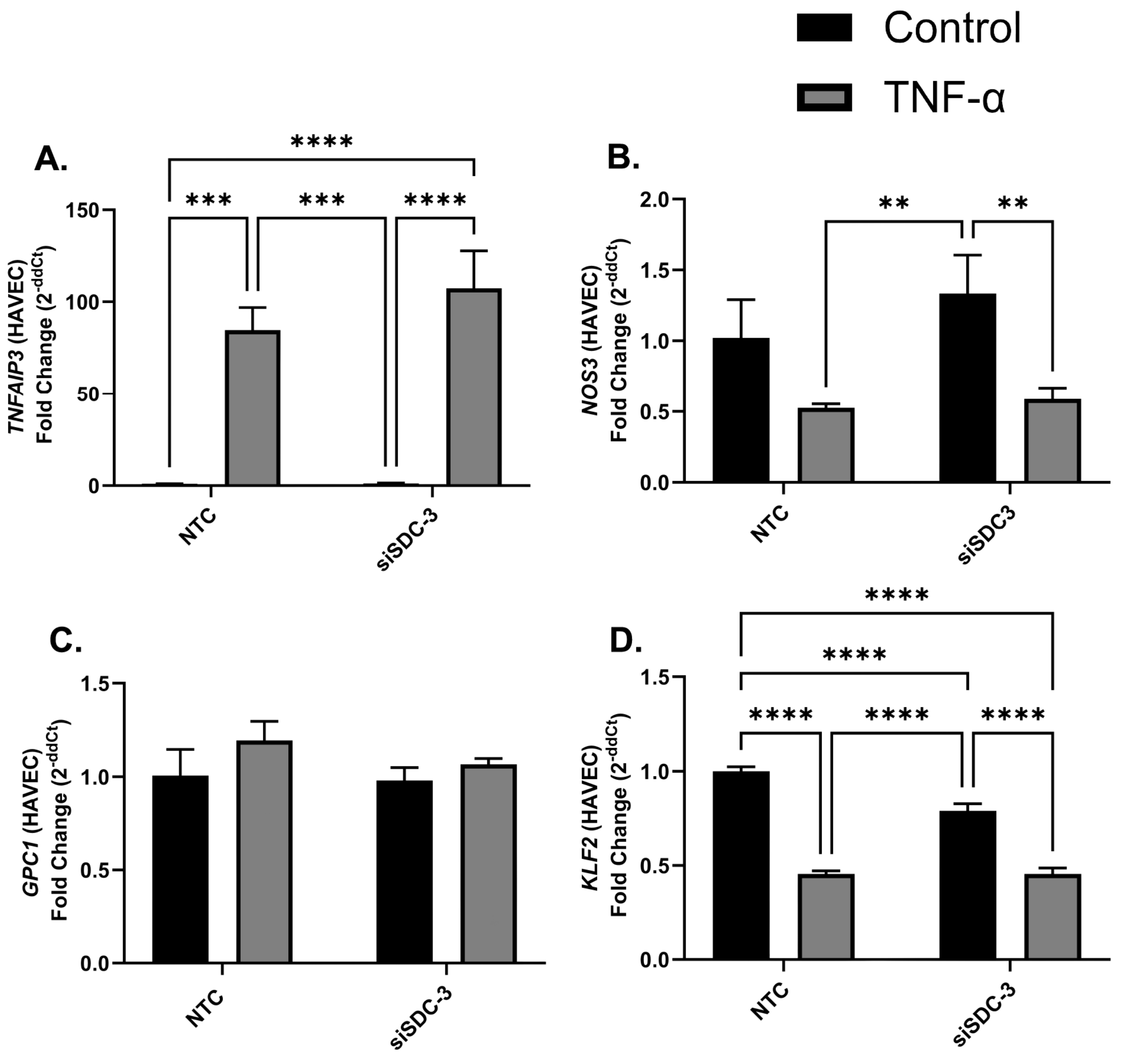


**Supplementary Figure 2. Effect of SDC3 knockdown on additional inflammatory and endothelial transcripts in HAVECs.** RT-qPCR quantification of (A) *TNFAIP3*, (B) *NOS3*, (C) *GPC1*, and (D) *KLF2* in NTC and siSDC3 HAVECs under control (black) or TNF-α–stimulated (100 ng/mL) (grey) conditions for 4 h, expressed as fold change relative to control (2^−ΔΔCt). Data are presented as mean ± SD (n = 3 per condition); *p < 0.05, **p < 0.01, ***p < 0.001, ****p < 0.0001.

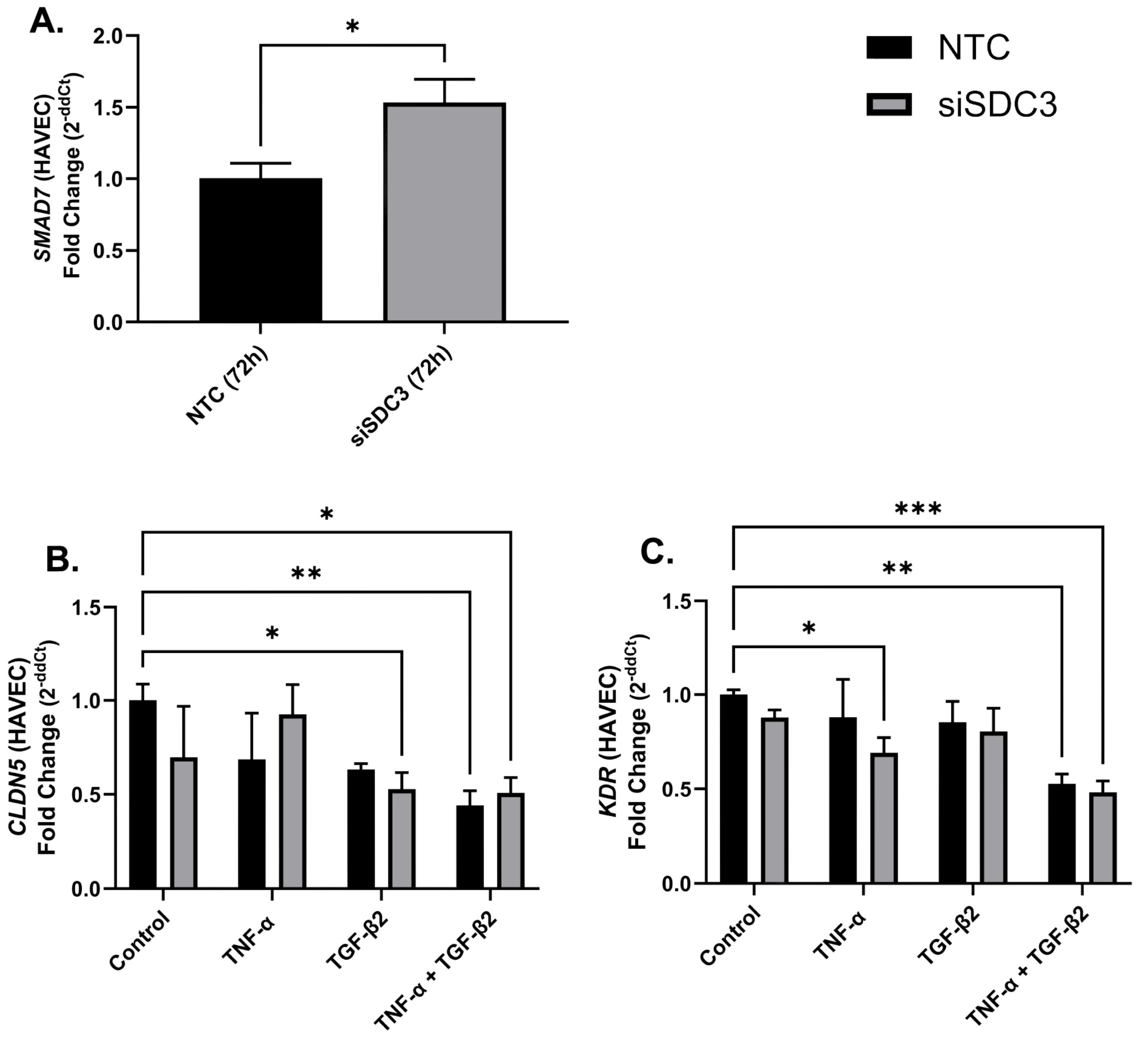


**Supplementary Figure 3. 72 Hour SMAD7 induction and endothelial marker expression in SDC3-silenced HAVECs.** (A) RT-qPCR quantification of *SMAD7* in NTC and siSDC3 HAVECs at 72 h post-transfection without stimulation, expressed as fold change relative to NTC (2^−ΔΔCt). (B, C) RT-qPCR quantification of the endothelial markers (B) *CLDN5* and (C) *KDR* in NTC (black) and siSDC3 (grey) HAVECs under control conditions or following stimulation with TNF-α, TGF-β2, or TNF-α + TGF-β2 (all 10 ng/mL) for 48 h, expressed as fold change relative to control (2^−ΔΔCt). Data are presented as mean ± SD (n = 2-3 per condition); *p < 0.05, **p < 0.01, ***p < 0.001, ****p < 0.0001.
